# Effects of Sanhuang plaster on expression of MyD88, TRAF6, MIP-1β and IL-23 in rats infected by MRSA

**DOI:** 10.1101/2022.03.02.482626

**Authors:** Haibang Pan, Qian Chen, Qi Fu, Tianming Wang, Xiaoli Li, Richeng Li, Mei Liu, Tiankang Guo

## Abstract

Objective: To investigate the effects of Sanhuang plaster on expression of myeloid differentiation factor 88 (MyD88), tumor necrosis factor receptor-related factor 6 (TRAF6), macrophage inflammatory protein-1β (MIP-1β) and its mediated cytokine interleukin-23 (IL-23) in soft tissues of rats infected by methicillin-resistant Staphylococcus aureus (MRSA). Methods: Ninety-six healthy rats were randomly divided into normal control group, model control group, Mupirocin group, high, medium and low dose groups of Sanhuang plaster, with 16 rats in each group. MRSA bacterial liquid was used to make skin and soft tissue infection models. The rats in the normal control group and the model control group were not given any treatment measures. The high, medium, and low dose groups of Sanhuang plaster were given to the affected area with Sanhuang plaster, and the Mupirocin group was given to the affected area for treatment for 10 days. The wound pathological changes were observed. The levels of MIP-1β and IL-23 in serum and infected tissues of rats in each group were measured by ELISA. The mRNA expressions of MyD88, TRAF6, MIP-1β and IL-23 were measured by Quantitative Real-time PCR, and Western blot was used to measure MyD88 and TRAF6 protein expression. Results: Compared with the model control group, the general condition of the Sanhuang plaster groups was significantly improved, and the pathological damage was reduced. The MIP-1β and IL-23 levels in the serum and infected tissues of the high dose group of Sanhuang plaster, the mRNA expressions of MyD88, TRAF6, MIP-1β and IL-23and in large and medium dose groups of Sanhuang plaster, and the protein expressions of Myd88 and TRAF6 in high dose of Sanhuang plaster were significantly decreased (p<0.05 or p<0.01). Conclusion: Sanhuang plaster may play a role in promoting healing of infected wounds by down-regulating the expression levels of MyD88, TRAF6, and MIP-1β, and inhibiting the abnormal secretion of cytokine IL-23.

## 1. Introduction

Sanhuang plaster is one of the unique hospital preparations in the affiliated Hospital of Gansu University of traditional Chinese Medicine. its main effects are promoting blood circulation and removing blood stasis, purging fire and detoxification, clearing heat and relieving pain, suppurating toxin and expelling pus, etc. it can be used to treat various infectious diseases such as furuncle, carbuncle, erysipelas, abscess and so on^[1]–[4]^. Although the clinical effect is remarkable, the mechanism of its anti-infection is still not clear. The purpose of this study was to further explore the mechanism of Sanhuang plaster in repairing wounds by observing the effects of Sanhuang plaster on the general survival status and pathological changes of infected tissues, MyD88, TRAF6, MIP-1 β and downstream cytokines IL-23 in MRSA infected model rats.

## 2. materials and methods

### 2.1 Materials

#### 2.1.1 Experimental animal

A total of 96 SD rats with body mass 170g-190g were purchased from the Medical Experimental Center of Gansu University of traditional Chinese Medicine, certificate number: SCXK (Gan) 2015-0002.Feeding environment: SPF animals are raised in the room, the humidity (45% Mel 55%) and room temperature (22°C-25°C) are controlled throughout the process.

#### 2.1.2 Main drugs, reagents and instruments

Sanhuang Plaster (affiliated Hospital of Gansu University of traditional Chinese Medicine, batch number: Z04010878); Mupirocin(Sino American Tianjin SmithKline Pharmaceutical Co., Ltd. batch number: 3L4K); GAPDH Antibodies(ImmunoWay inc); ELISA kits for MIP-1β and IL-23(Shanghai Future Industrial Co., Ltd.); Real-time fluorescence quantitative PCR kit and reverse transcription kit(TaKaRa Co., Ltd.); MyD88 Anti-body TRAF6 Anti-body MIP-1β Anti-body and IL-23 Anti-body (GeneTexCo., Ltd.); Horseradish enzyme labeled goat anti-rabbit IgG(Beijing Zhongshan Jinqiao Biotechnology Co., Ltd.); Albumin Bovine V, 30%Glue making liquid, Tris 6.8, PMSF, SDS, Glycine, Tris 8.8(Beijing Soleibao Technology Co., Ltd.); CT14RD high speed cryogenic centrifuge(Shanghai Tianmei biochemical instrument and equipment Engineering Co., Ltd.); SCIENTZ-48 High Throughput Tissue Grinder(Ningbo Xinzhi Biotechnology Co., Ltd.); BIOMATE 3s nucleic acid protein tester, Benchmark Plus enzyme-linked immunosorbent assay(American Bio-Rad Co., Ltd.).

### 2.2 Method

#### 2.2.1

Grouping and treatment of rats 96 SD rats were divided into 6 groups: normal control group, model control group, Mupirocin group, Sanhuang plaster high, medium and low dose groups, with 16 rats in each group.The model was established by referring to the method designed by NataliaMalachowa et al^[5]^. That is, 24 hours before the experiment, 8%Na_2_S was used to mark the area of 2cm×2cm with a signature pen on the back of each mouse near the cervical side with the spine as the midline. After hair removal and subcutaneous injection of 1×10^7^ CFU/mL MRSA solution determined by the pre-test, the general condition of rats and subcutaneous soft tissue infection were observed and recorded every day. All groups began to apply Sanhuang plaster on the next day according to the experimental design. High, medium and low dose Sanhuang plaster groups (1g, 0.5g and 0.25g/ of Sanhuang plaster with 1g/ of vaseline, the use dose is equivalent to 4 times, 2 times and 1 times with reference to the dose used in the corresponding clinical area), Mupirocin group (0.5g/ of Mupirocin with 1g/ of vaseline, the use dose is equivalent to 2 times with reference to the dose used in the corresponding clinical area), normal control group and model control group (1g/ of Vaseline), half male and half female, and raised in a single cage. The wound was cleaned with normal saline for the first time, and then was applied externally, and the dressing was changed every 12 hours.

#### 2.2.2

Histopathological changes of infected rats in each group, the junction tissue between wound and normal skin of rats in each group was fixed, embedded and sliced, and stained with HE dye.

#### 2.2.3

The contents of MIP-1β and IL-23 in serum and infection group were detected by ELISA method in accordance with the instructions of ELISA kit for follow-up operation. The OD value of each hole was determined by enzyme labeling instrument, and 450nm was selected as the measurement wavelength.

#### 2.2.4

The mRNA expressions of MyD88, TRAF6, MIP-1β and IL-23 in infected tissues were detected by real-time fluorescence quantitative PCR. The total RNA was extracted by Trizol and amplified after reverse transcription. The primer was synthesized by Bao Biological Engineering (Dalian) Co., Ltd. β-action is used as the internal reference gene, and the reaction system and parameters are set according to the instructions of GoTaq3-Step Real-time qPCR kit.

#### 2.2.5

The protein expression of MyD88 and TRAF6 in infected tissue was detected by Westernblot. Appropriate amount of tissue protein was frozen in refrigerator, tissue protein was extracted and quantified, SDS-PAGE electrophoresis was carried out, membrane was transferred, sealing solution was added, first antibody was added, shaker at 4°C overnight, sheep anti-rabbit second antibody was added after washing, and the expression of MyD88 and TRAF6 was calculated by Quantityone software after incubation at room temperature for 1 hour.

### 2.3 Statistical analysis

SPSS21.0 software was used for analysis. The experimental data were expressed by mean ± standard deviation 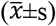. The analysis of variance was used for comparison between groups, the homogeneity of variance was compared with LSD method, and the variance was compared with Tanmhane’s T2 method. P <0.05 indicates that the difference is statistically significant.

## 3 Result

### 3.1 Effect of Sanhuang plaster on histopathological changes of infected rats in each group

The structure of skin and soft tissue of rats in normal control group was clear under microscope (figure 1:A). In the model group, the epidermis squamous epithelium was generally thickened, the epidermis structure near the abscess was destroyed, the hierarchy was unclear, and a large number of inflammatory cells infiltrated; the dermis became thinner and the staining deepened, and the collagen fibers coagulated in flakes, lumps, strips, accompanied by a large number of inflammatory cell infiltration, and the subcutaneous tissue structure decreased (figure 1: B). In the Mupirocin group, the hierarchical structure of the epidermis of the infected tissue was unclear; a large number of pus cells and neutrophils could be seen in the necrotic tissue; the dermis became thinner and the staining deepened, and the collagen fibers were flake, lump, cord-like and accompanied with a large number of inflammatory cell infiltration; the subcutaneous tissue structure decreased (figure 1:C). In the high-dose Sanhuang plaster group, the granulation tissue around the infected tissue abscess was filled with thin-walled capillaries (figure 1: D). The degree of infection tissue injury in the middle and low dose groups was more serious, the necrotic layer was higher than that in the high dose group, and the filling of granulation tissue around it was less than that in the high dose group (figure 1: E, F).

**Figure 1.**
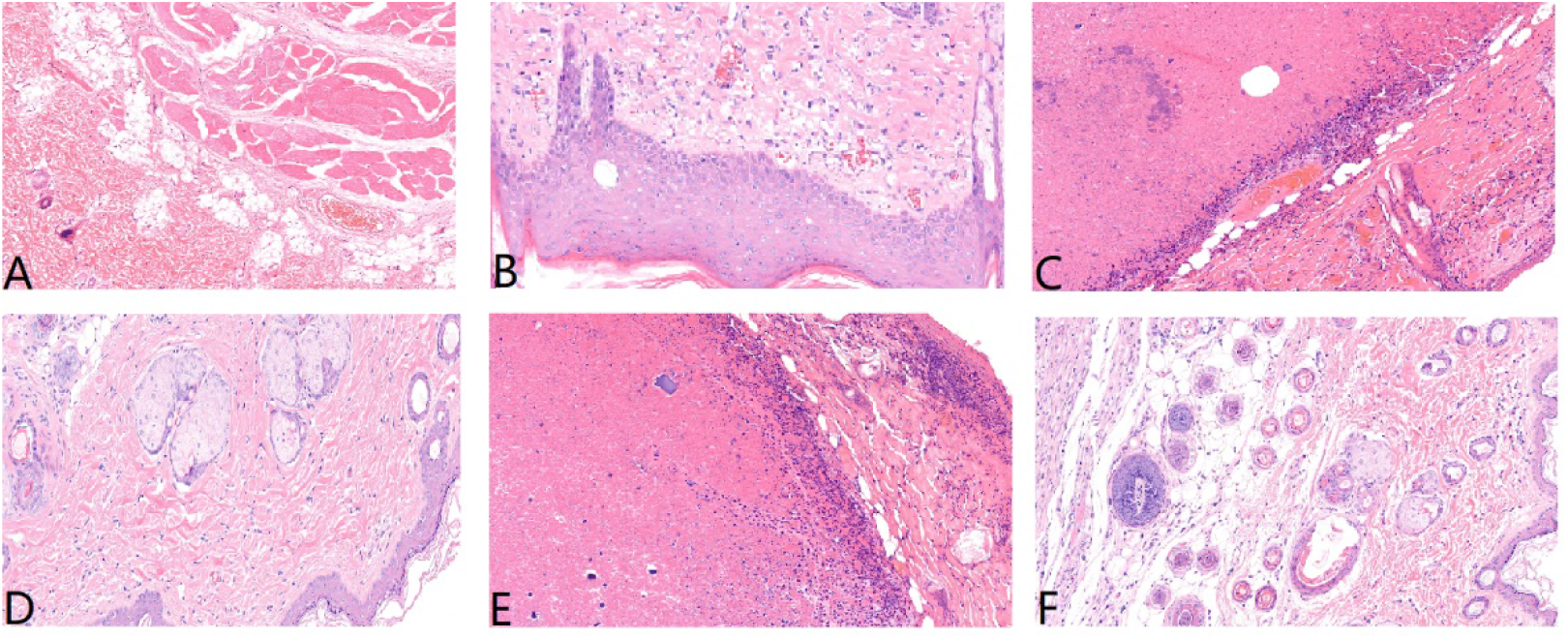
Effect of Sanhuang Plaster on pathological changes of MRSA infected rats (HE, ×400) A.Normal control; B.Model group; C.Mupirocin group; D.Sanhuang plaster high dose group; E.Sanhuang plaster medium dose group; F.Sanhuang plaster low dose group

### 3.2 Effect of Sanhuang plaster on the contents of MIP-1β and IL-23 in serum and infected tissues of rats in each group

Compared with the normal control group, the contents of MIP-1β and IL-23 in the serum and infected tissue of the model control group increased significantly, while the contents of MIP-1β and IL-23 in the serum and infected tissue decreased significantly in the Mupirocin group and Sanhuang plaster high dose group (p<0.05), but not in the low dose group (p<0.05). The results are shown in Table 1.

**Table 1.**
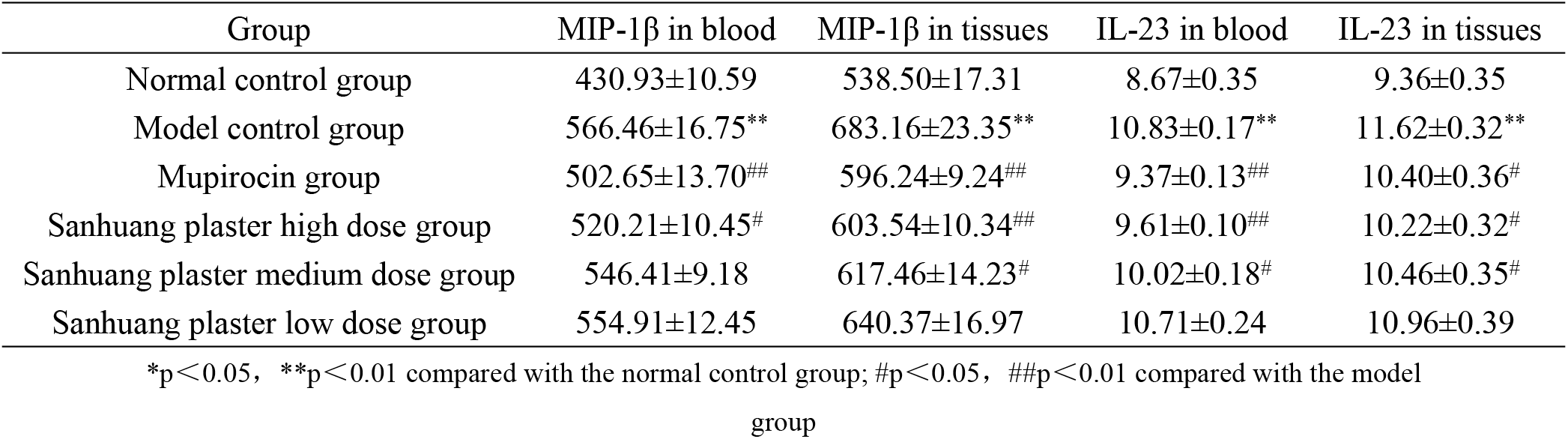
Effect of Sanhuang plaster on the contents of MIP-1β and IL-23 in serum and infected tissues of rats in each group(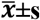,n=16)

### 3.3 Effect of Sanhuang plaster on the levels of MyD88, TRAF6, MIP-1β and IL-23 mRNA in infected tissues of rats in each group

Compared with the normal control group, the mRNA expression levels of MyD88, TRAF6, MIP-1 β and IL-23 in the infected tissue of the model control group were significantly increased (p < 0.01). Compared with the model control group, the mRNA expression levels of MyD88, TRAF6, MIP-1 β and IL-23 in infected tissues of rats in Mupirocin group and Sanhuang plaster high dose group were significantly lower than those in model control group (p<0.05).There was no significant decrease in low dose group (p>0.05). The results are shown in Table 2.

**Table 2.** Effect of Sanhuang plaster on the levels of MyD88, TRAF6, MIP-1β and IL-23 mRNA in infected tissues of rats in each group (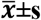,n=16)

| Group | Myd88 | TRAF6 | MIP-1β | IL-23 |
| --- | --- | --- | --- | --- |
| Normal control group | 1.01±0.05 | 1.01±0.05 | 1.00±0.03 | 1.00±0.06 |
| Model control group | 1.71±0.05** | 1.65±0.06** | 1.49±0.04** | 1.56±0.05** |
| Mupirocin group | 1.48±0.06<sup>#</sup> | 1.13±0.13<sup>##</sup> | 1.24±0.05<sup>##</sup> | 1.21±0.14<sup>##</sup> |
| Sanhuang plaster high dose group | 1.19±0.04<sup>##</sup> | 1.28±0.06<sup>##</sup> | 1.29±0.04<sup>##</sup> | 1.33±0.08<sup>#</sup> |
| Sanhuang plaster medium dose group | 1.45±0.05<sup>##</sup> | 1.51±0.09 | 1.34±0.03<sup>#</sup> | 1.47±0.03 |
| Sanhuang plaster low dose group | 1.49±0.06 | 1.53±0.09 | 1.44±0.05 | 1.52±0.16 |

\*p<0.05, \*\*p<0.01 compared with the normal control group; #p<0.05, ##p<0.01 compared with the model group

### 3.4 Effect of Sanhuang plaster on the contents of MyD88 and TRAF6 protein in infected tissues of rats in each group

Compared with the normal group, the contents of Myd88 and TRAF6 protein in infected tissues of the model control group increased significantly, while the contents of MyD88 and TRAF6 protein in infected tissues decreased significantly in the Mupirocin group and Sanhuang plaster high dose group (p <0.05), but not in the middle and low dose groups (p >0.05). The results are shown in Table 3 and figure 2.

**Figure 2.**
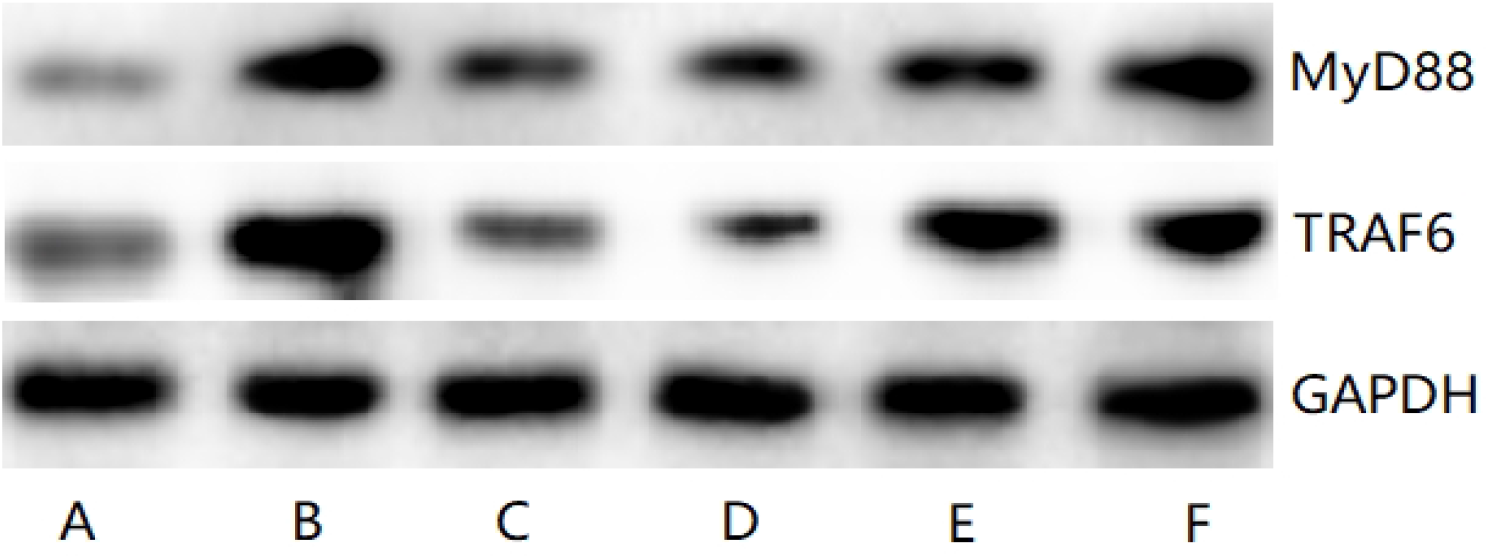
Western blotting of MyD88 and TRAF6 in infected tissues of rats in each group. A.Normal control; B.Model group; C.Mupirocin group; D.Sanhuang plaster high dose group; E.Sanhuang plaster medium dose group; F.Sanhuang plaster low dose group

**Table 3.** Effect of Sanhuang plaster on the contents of MyD88 and TRAF6 protein in infected tissues of rats in each group (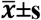,n=16)

| Group | MyD88 | TRAF6 |
| --- | --- | --- |
| Normal control group | 0.50±0.02 | 0.64±0.04 |
| Model control group | 0.83±0.07** | 1.58±0.35* |
| Mupirocin group | 0.63±0.03<sup>##</sup> | 0.71±0.04<sup>#</sup> |
| Sanhuang plaster high dose group | 0.59±0.04<sup>##</sup> | 0.79±0.06<sup>#</sup> |
| Sanhuang plaster medium dose group | 0.73±0.05 | 1.17±0.15 |
| Sanhuang plaster low dose group | 0.91±0.10 | 1.34±0.26 |

\*p<0.05, \*\*p<0.01 compared with the normal control group; #p<0.05, ##p<0.01 compared with the model group

## 4. Discussion

Skin and Soft Tissue Infections (SSTI) is mainly an inflammatory reaction caused by pathogen invasion and reproduction, which often occurs in the course of infection in many clinical and surgical diseases. It can also enter the blood circulation and cause systemic suppurative infection ^[6][8]^. One of the most common pathogens in SSTI is Staphylococcus aureus, of which MRSA is the most common drug-resistant Staphylococcus aureus^[9][11]^. The previous study of our group showed that Sanhuang plaster has a good clinical effect on SSTI, in order to further explore the pathogenesis and treatment of SSTI, the rat model of SSTI was made by MRSA bacterial liquid, and the gene and protein expression levels of Myd88, TRAF6, MIP-1β and IL-23 in TLRs signal pathway were used as research indexes to study the mechanism of Sanhuang plaster in repairing wound.

Studies have shown that TLR2/4 signal pathway can activate the persistence of inflammatory response, strongly induce a “cascade” increase in the number of inflammatory factors, and then magnify the inflammatory response. TLRs signal pathway belongs to MyD88-dependent signal pathway, and its downstream factors can be activated by TRAF6, IL-1,Interleukin-1 receptor-associated kinase (IRAK1/4) and so on^[12][14]^. TRAF6 is one of seven closely related TRAF proteins, which can induce the activation of multiple kinase pathways. It is necessary for TLR superfamily signal transduction pathways^[15][17]^. Some studies pointed out that TRAF6 is the downstream effector protein of TLRs signal pathway and can regulate the release of inflammatory factors in this pathway^[18]^. The results showed that the contents of MyD88 and TRAF6 in infected tissues of the model control group were significantly increased, while the contents of MyD88 and TRAF6 were significantly decreased in the high dose group of Sanhuang plaster.

MIP-1 is a protein duplex corresponding to inflammatory activity. because of its inflammatory characteristics in vitro and in vivo, this protein mixture is named “macrophage inflammatory protein-1”. Its further differentiation produces two distinct but highly related proteins MIP-1α and MIP-1β, which are chemokine essential for the immune response of infection and inflammation^[19][20]^. MIP-1β is inducible in most mature hematopoietic cells and plays a major role in recruiting leukocytes to the infected site. In addition, chemokines often activate these cells, resulting in an increased local inflammatory response^[21][22]^. IL-23 is a heterodimer cytokine that acts as a proinflammatory cytokine^[23]^. In this study, the contents of MIP-1β and IL-23 in serum and infected tissues of rats in the model control group increased significantly, while those in the high dose group of Sanhuang plaster significantly decreased the contents of MIP-1β and IL-23.

To sum up, Sanhuang plaster can promote the healing of infected wound, and its mechanism may be related to the inhibition of TLRs signal pathway, down-regulation of MyD88 and TRAF6 gene and protein expression, inhibition of abnormal secretion of MIP-1β and pro-inflammatory factor IL-23 at the end of this pathway, and reduction of inflammatory cascade amplification.

